# TDP-43 dysfunction restricts dendritic complexity by inhibiting CREB activation and altering gene expression

**DOI:** 10.1101/2019.12.12.874735

**Authors:** Josiah J. Herzog, Mugdha Deshpande, Weijin Xu, Reazur Rahman, Hannah Suib, Michael Rosbash, Avital A. Rodal, Suzanne Paradis

**Author notes:** Equal Contribution (Josiah J. Herzog, Mugdha Deshpande).

## Abstract

Amyotrophic lateral sclerosis (ALS) and frontotemporal dementia (FTD) are two related neurodegenerative diseases that present with similar TDP-43 pathology in patient tissue. TDP-43 is an RNA-binding protein and forms aggregates in neurons of ALS and FTD patients as well as in a subset of patients diagnosed with other neurodegenerative diseases. Despite our understanding that TDP-43 is essential for many aspects of RNA metabolism, it remains obscure how TDP-43 dysfunction contributes to neurodegeneration. Interestingly, several neurological disorders display altered dendritic morphology and complexity, which are thought to precede neurodegeneration. In this study, we used TRIBE (targets of RNA-binding proteins identified by editing) as a new approach to identify signaling pathways that regulate dendritic branching downstream of TDP-43. We found that TDP-43 targets are enriched for pathways that signal to the CREB transcription factor. We further found that TDP-43 dysfunction inhibits CREB activation and CREB transcriptional output, and restoring CREB signaling rescued defects in dendritic branching. Our data therefore provide a novel mechanism by which TDP-43 dysfunction interferes with dendritic branching, and define new pathways for therapeutic intervention in neurodegenerative diseases.

## Introduction

Amyotrophic lateral sclerosis (ALS) is a debilitating and rapidly progressing neurodegenerative disease that is often co-morbid with Frontotemporal Dementia (FTD). A dominantly heritable form of ALS/FTD is caused by mutations in the gene TARDBP, which encodes for the TDP-43 protein (Sreedharan et al. 2008; Kovacs et al. 2009). TDP-43 is also a major component of pathological protein inclusions in more than 95% of sporadic ALS and ∼45% of FTD, suggesting that it is a central underlying factor in disease (Arai et al. 2006; Neumann et al. 2006; Ling et al. 2013). TDP-43 is an RNA-binding protein that shuttles between the nucleus and cytoplasm, and has been implicated in a variety of cellular functions including DNA transcription, RNA splicing, miRNA processing, RNA transport and translation (Nussbacher et al. 2019; Prasad et al. 2019). Upon cellular insult, TDP-43 redistributes to the cytoplasm, where it associates with liquid stress granules, and can further irreversibly aggregate under pathological conditions (Birsa et al. 2019; Prasad et al. 2019). Importantly, this TDP-43 cytoplasmic accumulation results in nuclear depletion of endogenous TDP-43, leading to the proposal that toxic gain of function of TDP-43 in the cytoplasm with concomitant TDP-43 loss of function in the nucleus contributes to the disease (Xu 2012; Birsa et al. 2019).

We previously reported that both increased or decreased TDP-43 expression (modeling gain and loss of function, respectively) results in reduced dendritic outgrowth and complexity without an immediate effect on cell viability (Herzog et al. 2017). In vivo, a major consequence of altered dendritic elaboration is disruption to overall neuronal circuit connectivity and cell-to-cell communication (Lefebvre et al. 2015; Stuart and Spruston 2015), which has been suggested to be an underlying feature of the early stages of ALS and FTD (Kulkarni and Firestein 2012; Lopez-Domenech et al. 2016; Kweon et al. 2017). Importantly, the decreased dendritic branching observed upon TDP-43 overexpression is dependent on the RNA-binding ability of the protein (Herzog et al. 2017). However, it remains unclear which TDP-43 RNA targets impinge on many potential extrinsic and intrinsic signaling pathways that promote proper dendrite elaboration (Dong et al. 2015).

One obvious way forward is to identify TDP-43 RNA targets relevant to neurodegeneration (Polymenidou et al. 2011; Sephton et al. 2011; Tollervey et al. 2011). Using <u>R</u>NA <u>i</u>mmuno<u>p</u>recipitation followed by deep sequencing (RIP-seq) and <u>c</u>ross-<u>l</u>inking <u>i</u>mmuno<u>p</u>recipitation coupled with high-throughput sequencing (CLIP-seq and its variants), other groups independently generated lists of several thousand potential TDP-43 targets in the mammalian brain. Because of this large number and because RIP and CLIP have potential limitations (Wheeler et al. 2018), we applied an orthogonal approach to identify TDP-43 target RNAs.

TRIBE (<u>T</u>argets of <u>R</u>NA-binding proteins <u>I</u>dentified <u>B</u>y <u>E</u>diting) is an antibody-independent method to identify RBP target mRNAs (McMahon et al. 2016; Xu et al. 2018), which had only been previously applied to the *Drosophila* system. To identify putative TDP-43-bound target RNAs, we performed TRIBE in RNA obtained from primary cultures of mammalian neurons in which TDP-43 expression was altered. From the RNA target list generated, we identified multiple RNAs encoding various signaling pathway components. They including those focused on signaling via the activity-regulated cAMP response element-binding protein (CREB) transcription factor (Alberini 2009).

Transcriptional programs such as those downstream of CREB activation control the elaboration of neuronal dendritic arbors (Dong et al. 2015). Upstream regulation of CREB activity is complex and involves multiple upstream signal transduction networks including the CaMK, MAPK and PKA pathways (Tan et al. 1996; Tao et al. 1998; Watson et al. 2001; Redmond et al. 2002; Cohen and Greenberg 2008; Alboni et al. 2011). Some of these pathways contain TDP-43 targets, and we validated that the CREB transcriptional activity is misregulated upon TDP-43 dysfunction, leading in turn to defects in dendrite morphogenesis. Restoring CREB activity restores dendrite elaboration, suggesting that this pathway may be a new avenue for therapeutic intervention in TDP-43-related neurological diseases.

## Results

### TDP-43 RNA targets are involved in multiple signaling pathways that regulate the CREB transcription factor

TDP-43 is an RBP that has been suggested to interact with several thousand RNA transcripts in neuronal cells (Polymenidou et al. 2011; Sephton et al. 2011; Tollervey et al. 2011). RIP or CLIP methods were used in these studies, but they have potential liabilities, e.g., reassociation of RNAs to RBPs after cell lysis (Mili and Steitz 2004) and poor efficiency of cross-linking (Darnell 2010), respectively. These approaches also rely heavily on high-quality antibodies, which varied between these studies. These limitations make the identification of true target genes vs false positive and false negatives difficult to assess. Not surprisingly perhaps, the lists of putative target transcripts do not correlate well with each other, leading to considerable uncertainty about TDP-43 targets. An orthogonal approach like TRIBE is independent of antibodies and ideal for contributing to identifying bona fide TDP-43 targets.

The original TRIBE method takes advantage of the *Drosophila* RNA editing enzyme ADAR (referred to henceforth as dADAR), which endogenously interacts with specific structured RNAs and converts adenosine to inosine. Fusion of the ADAR catalytic domain (ADARcd) to an RBP of interest and expression *in vivo* directs the fusion protein to target RNAs of that RBP, on which the ADARcd then performs A-to-I editing. These editing events are detected as A-to-G mutations through standard RNA-seq and computational analysis (McMahon et al. 2016; Xu et al. 2018). To adapt the TRIBE method to mammalian systems, we used the catalytic domain of human ADAR2 (hADARcd), the human homolog of dADARcd, rather than dADARcd.

A pilot experiment was first performed in HEK293T cells to test the effectiveness of the TRIBE system in mammalian cells. Expression of the TDP-43 TRIBE fusion protein enhanced the number of editing sites by ∼150 times (Fig. 1A). Moreover, no above-background editing was detected in cells expressing TDP43-5FL TRIBE, a non-RNA-binding mutant of TDP-43 (Herzog et al. 2017), or the hADARcd alone (Fig. 1A). The data not surprisingly show that the efficiency and specificity of mammalian TRIBE is dependent on the RBP as previously indicated from the *Drosophila* system (McMahon et al. 2016; Xu et al. 2018).

**Figure 1:**
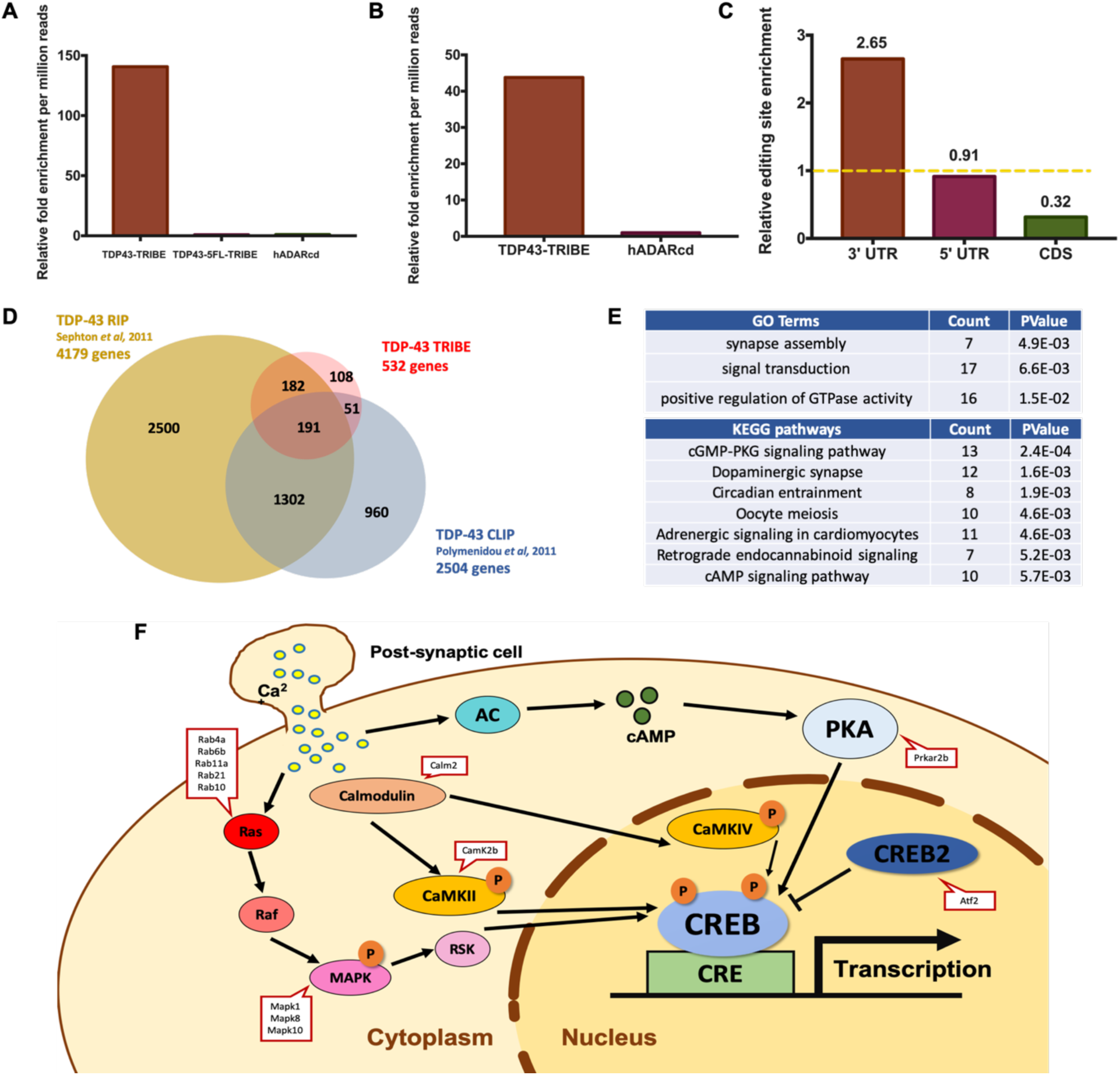
TDP-43 TRIBE reveals TDP-43 RNA targets involved in CREB signaling. **(A)** Quantification of the relative number of editing sites detected in HEK293T cells transfected with plasmids expressing TDP-43 TRIBE, TDP-43-5FL TRIBE or hADARcd expressed under control of the CMV promoter (N=2). All the editing sites shown here and in all figures were present in both replicates. Y-axis is the relative fold change to hADARcd and normalized by their sequencing depth. A requirement of 10 reads of 5 percent A-to-G editing is imposed to qualify as an editing site. (See method section for details) **(B)** Quantification of the relative number of editing sites detected in cultured rat cortical neurons transfected with lentivirus expressing TDP-43 TRIBE or hADARcd under control of the human synapsin promoter (N=2). Normalization performed as in (A). **(C)** Quantification of TDP-43 TRIBE editing sites enrichment in each mRNA region. Editing sites in 3’UTR, 5’UTR or CDS were assigned based on RefSeq annotation, and enrichment was normalized to sequencing coverage. For example, 76% of all editing sites were located in 3’UTR, but 3’UTR only accounts for 29% of all sequencing depth, so it is 2.65-fold enriched. **(D)** Venn diagram demonstrating the 191 genes that overlap between 3 different studies, including ours, that sought to identify TDP-43 target RNAs. TDP-43 TRIBE identified 108 unique genes that were not identified previously by RIP-seq or CLIP-seq. **(E)** Top GO-term and KEGG pathway hits with DAVID functional annotation analysis performed on TDP-43 TRIBE target list. The gene list was normalized to the background of the top 3000 expressed genes in cultured rat cortical neurons (see methods). **(F)** Diagram of key signaling pathways related to CREB activation. TDP-43 TRIBE target genes are listed in red squares.

To identify RNA targets of TDP-43 in the central nervous system, we performed TDP-43 TRIBE in cultured rat cortical neurons, a frequently used system for studying the neurodegenerative pathology of TDP-43 (Herzog et al. 2017; Baskaran et al. 2018). We constructed lentiviral vectors encoding TDP-43 TRIBE or the hADARcd alone under the human synapsin 1 promoter (hSyn). After viral packaging, we transduced the viruses into cultured cortical neurons at 4 days in vitro (DIV). At 7 DIV, we harvested RNA, made mRNA-seq libraries and performed high-throughput sequencing. Analysis of the sequence data with our computational pipeline (Rahman et al. 2018) identified 532 genes, hosting 1162 editing sites, as TDP-43 targets; this was ∼45 fold more editing sites than the hADARcd control (Fig. 1B, 1D). Analysis of the editing distribution within each mRNA region revealed a dramatic enrichment in 3’ UTR regions (Fig. 1C), suggesting a strong preference of TDP-43 for 3’ UTR binding. This general 3’ UTR preference agrees with a previous publication (Tollervey et al. 2011) and has also been described for a handful of specific mRNAs (Polymenidou et al. 2011; Sun et al. 2014; Fukushima et al. 2019). 3’ UTR binding also coincides with the proposed role of TDP-43 as an RNA stability and translational regulator (Tank et al. 2018; Neelagandan et al. 2019).

To examine how well the TDP-43 TRIBE target identification agrees with the RIP data (Sephton et al. 2011) and CLIP data (Polymenidou et al. 2011), we compared the target lists from the three datasets. Due to unavailability of the full gene list from Polymenidou et al., we downloaded these raw data via the GEO database and processed them using updated RNA-seq analysis standards, which reduced the number of target genes from 6304 to 2504.

TDP-43 TRIBE identified fewer targets than the other two methods. This is most simply explained by a combination of a lower false-positive rate and higher false-negative rate with TRIBE. Another possibility is the absence of edited intronic sequences in our data, i.e., nuclear targets, due to use of pA RNA. (See Discussion.) In any case, 191 high-confidence genes were identified in common between all 3 lists (Fig. 1D). This is a very large and highly significant fraction of the number of identified TRIBE genes. Interestingly and despite the large difference in the number of target genes, the TRIBE data agree better with the RIP data (∼70% of TRIBE targets are RIP targets) than with the CLIP data (Fig. 1D, Z-test performed, p<0.001). The results taken together suggest that TDP-43 TRIBE is a viable method to identify RBP RNA targets in mammalian cells.

To gain insight into the biological processes potentially impacted by TDP-43, we used DAVID functional annotation tools for Gene Ontology term (GO term) and KEGG pathway analysis of TDP-43 TRIBE targets. From our TDP-43 TRIBE target list of 532 genes, both GO and KEGG generated lists of pathways and processes relevant to neuronal signaling and synapses (Fig. 1E), consistent with our phenotypic observations (Herzog et al. 2017). The results were similar when the 191 high-confidence genes were fed into the same analysis (Table S1). These findings suggest that the shorter TDP-43 TRIBE target list more likely represents bona fide TDP-43 targets.

More specifically, the genes identified by TDP-43 TRIBE are enriched for GO terms involving dendritic branching including synapse assembly. These genes are also enriched in KEGG pathways that signal to the CREB transcription factor (Fig. 1E, 1F). CREB is involved in dendritic elaboration and lies downstream of several signaling stimuli (Tan et al. 1996; Tao et al. 1998; Watson et al. 2001; Redmond et al. 2002; Alboni et al. 2011). They result in CREB activation through phosphorylation at serine 133 (pCREB), which together with recruited coactivators activates downstream gene expression (Shaywitz and Greenberg 1999). Notably, genes from TDP-43 TRIBE are enriched for pathways including cAMP, Hippo, MAPK, Adrenergic signaling, and cGMP-PKG (Figure 1E), which result in CREB phosporylation. Specifically, TDP-43 TRIBE targets include CaMKII, Calmodulin, Ras family members, MAPKs, PKA and CREB2 (Figure 1F), suggesting that CREB transcriptional regulation plays a central role in TDP-43-dependent neuronal function.

### TDP-43 dysfunction inhibits CREB activation

We next sought to determine if TDP-43 dysfunction affects CREB activation in cultured mammalian neurons. We isolated cortical neurons from E18 rat pups, cocultured the neurons on a glial monolayer, and transfected them at 2 DIV with a vector expressing GFP and an empty vector control, GFP and TDP-43 (i.e., TDP-43 OE), or GFP and a shRNA targeting TDP-43 (i.e., TDP-43 KD). After 7 DIV, neurons were fixed and immunostained using either an antibody that specifically recognizes phosphorylated CREB at its canonical serine 133 activation site (pCREB) or total CREB protein. Relative levels of pCREB and CREB were calculated by normalizing signal intensity to endogenous TFIIS, a ubiquitous transcription elongation factor (Wind and Reines 2000).

We observed a significant decrease in pCREB signal in neurons that overexpress TDP-43 compared to controls (Fig. 2A, 2B). Similarly, pCREB intensity is decreased in neurons in which TDP-43 was knocked down (Fig. 2A, 2B). These results are consistent with out previous observation that dendritic branching is decreased by either TDP-43 OE or TDP-43 KD (Herzog et al. 2017). Total CREB levels remained constant in both TDP-43 OE and TDP-43 KD conditions compared to control (Fig. 2C, 2D), suggesting that TDP-43 dysfunction specifically inhibits CREB activation without affecting overall CREB protein abundance.

**Figure 2:**
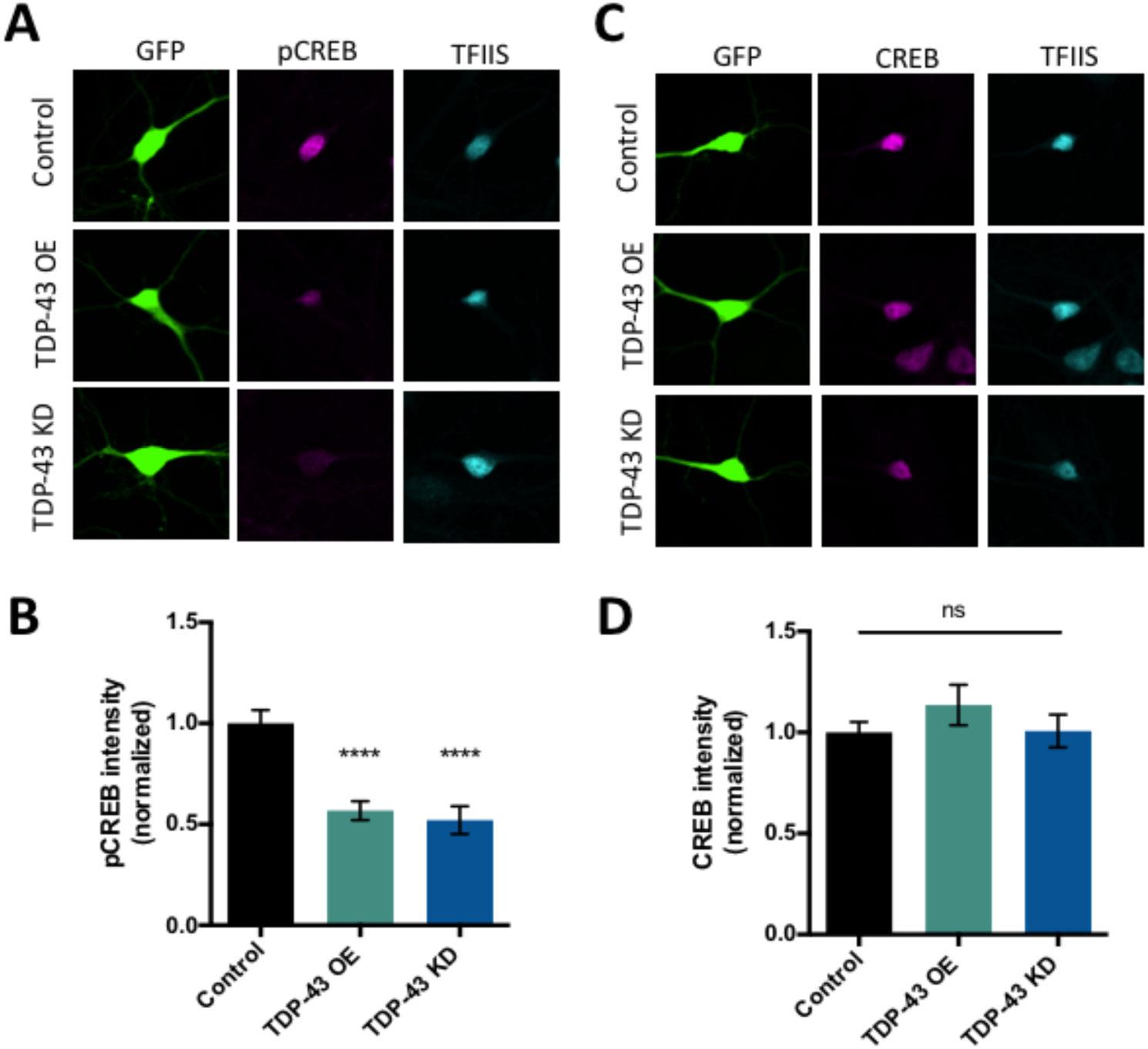
CREB phosphorylation is decreased by TDP-43 knock-down and overexpression. **(A)** Representative images of 7 DIV cultured cortical neurons transfected with plasmids expressing GFP and either empty vector control, TDP-43, or an shRNA targeting TDP-43 for knockdown immunostained with an antibody that recognizes phosphorylated CREB at Ser133 (pCREB). **(B)** Quantification of average fluorescence intensity for pCREB immunostaining normalized to the fluorescence intensity of the same neuron immunostained with an antibody that recognizes TFIIS. Control, N = 68; TDP-43 OE, N = 69, TDP-43 KD, N = 69. **(C)** Representative images of 7 DIV cultured cortical neurons transfected with plasmids expressing GFP and either empty vector control, TDP-43, or an shRNA targeting TDP-43 for knockdown immunostained with an antibody that recognizes total CREB protein. **(D)** Quantification of average fluorescence intensity for CREB immunostaining normalized to the fluorescence intensity of the same neuron immunostained with an antibody that recognizes TFIIS. Control, N = 70; TDP-43 OE, N = 68, TDP-43 KD, N = 65. **** p< 0.0001, one-way ANOVA, error bars represent standard error of the mean.

### CREB transcriptional output is suppressed by TDP-43 dysfunction

One likely consequence of the effect of TDP-43 dysfunction on CREB phosphorylation is suppression of CREB-dependent transcriptional activity. Therefore, we investigated whether TDP-43 dysfunction interfered with activity-dependent CREB transcriptional output using a dual luciferase assay to quantify CREB transcriptional activity. To this end, we transfected primary cultures of rat cortical neurons at 2 DIV with either an empty vector control, a plasmid expressing (TDP-43 OE), or an shRNA targeting TDP-43 (TDP-43 KD) along with a reporter construct containing 4 CREB response elements (4xCRE) upstream of a firefly luciferase gene and a control reporter expressing Renilla luciferase from the thymidine kinase promoter. At 7 DIV, a subset of control, TDP-43 OE, and TDP-43 KD neuronal cultures were treated with 55 mM KCl for 6 hours to mimic neuronal depoloarizaton and induce CREB activation (Tao et al. 2002). After 6 hours, cell lysates were harvested and firefly and Renilla luminescence was quantified for each condition. We observed a significant decrease in KCl-dependent CREB transcriptional output upon either TDP-43 OE or TDP-43 KD (Fig. 3). Taken together, our data suggests that TDP-43 dysfunction inhibits CREB activation and suppresses CREB transcriptional activity.

**Figure 3:**
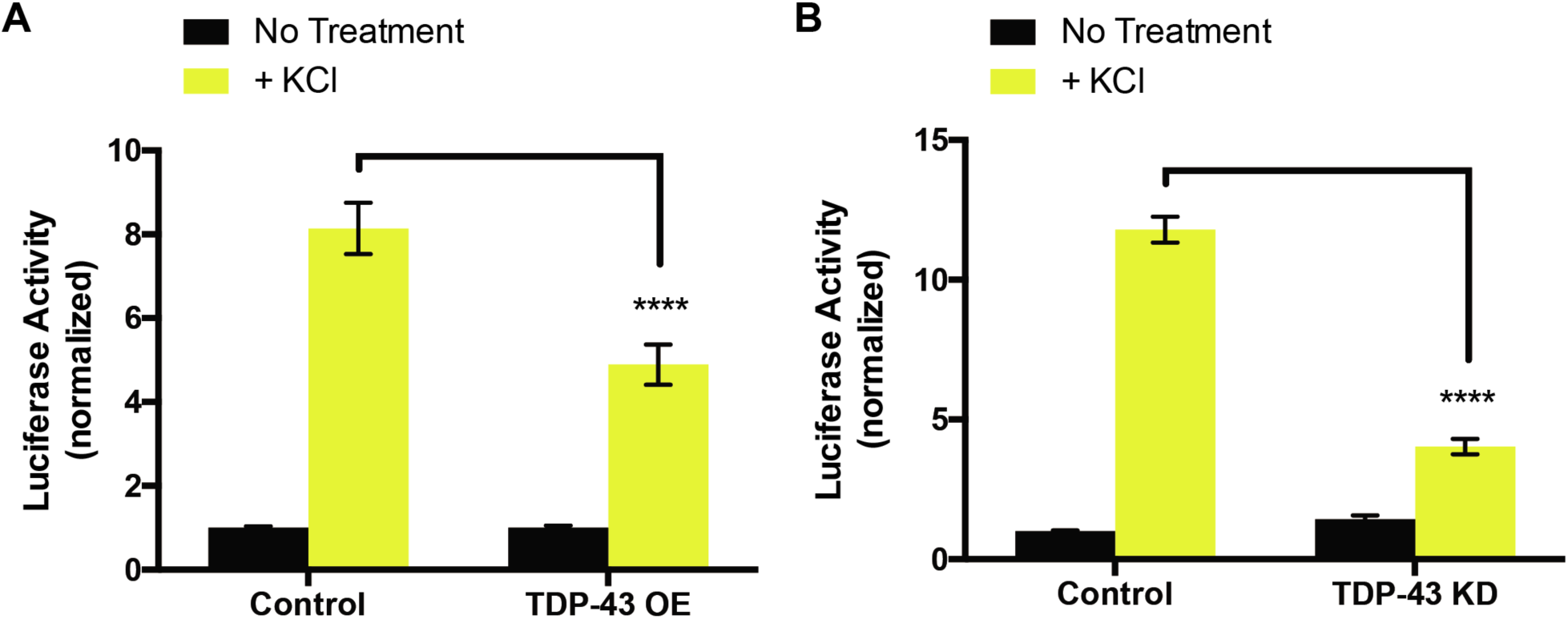
KCl-dependent CREB transcriptional activity is suppressed by TDP-43 dysfunction. Quantification of luciferase activity (firefly luciferase luminescence/Renilla luciferase luminescence) from cell lysates of cortical neurons transfected with an empty vector control, (A) TDP-43, or (B) an shRNA targeting TDP-43 for knockdown. Yellow bars represent conditions where a subset of cortical neurons were exposed to 55 mM KCl for 6 hours and black bars represent cortical neurons that were not stimulated with KCl. To account for changes in transfection efficiency, Firefly was normalized to Renilla for each condition. Each condition was normalized to the “No treatment” control condition for each experiment. Each experiment was performed in triplicate and repeated at least 3 times. Data is presented as the mean of normalized luciferase activity. Two-way ANOVA with Tukey’s test. **** = p < 0.0001. Error bars represent standard error of the mean.

### Activation of CaMK pathways restores dendritic branching defects induced by TDP-43 dysfunction

Calcium/calmodulin-dependent protein kinase IV (CaMKIV) stimulates dendritic growth via CREB phosphorylation at Ser133 and subsequent activation (Redmond et al. 2002; Ghiretti et al. 2013). We therefore asked if expression of a constitutively active form of CaMKIV (CaMKIV CA) (Wayman et al. 2006; Wayman et al. 2008) would restore dendritic branching under conditions in which TDP-43 expression is increased or decreased. CaMKIV was made constitutively active by fusing a nuclear localization signal; expression of this construct in hippocampal neurons has previously been shown to increase dendritic complexity (Ghiretti et al. 2013). Hippocampal neurons isolated from E18 rat pups were cocultured on a glial monolayer and transfected at 2 DIV with GFP and either an empty vector control, a plasmid expressing CaMKIV CA alone or in combination with appropriate constructs to achieve TDP-43 OE or TDP-43 KD (as described above). At 5 DIV, these neurons were fixed, imaged, and analyzed for dendritic complexity by Sholl Analysis (Sholl 1953). Expression of CaMKIV CA alone increased dendritic complexity above control levels, consistent with previous reports (Redmond et al. 2002; Ghiretti et al. 2013). We also observed decreased dendritic branching in both TDP-43 OE and TDP-43 KD conditions similar to previous reports (Figure 4A-C) (Majumder et al. 2012; Schwenk et al. 2016; Herzog et al. 2017). Our data demonstrate that expression of CaMKIV CA in neurons in which TDP-43 has been knocked down partially restored the reduction in dendritic branching (Fig. 4C). In contrast, we found that expression of CaMKIV CA in neurons overexpressing TDP-43 did not have a significant effect on branching. The fact that boosting CaMK signaling compensates for lack of TDP-43, but not TDP-43 overexpression, suggests that there are notable differences in upstream signaling pathways that are impacted by these manipulations.

**Figure 4:**
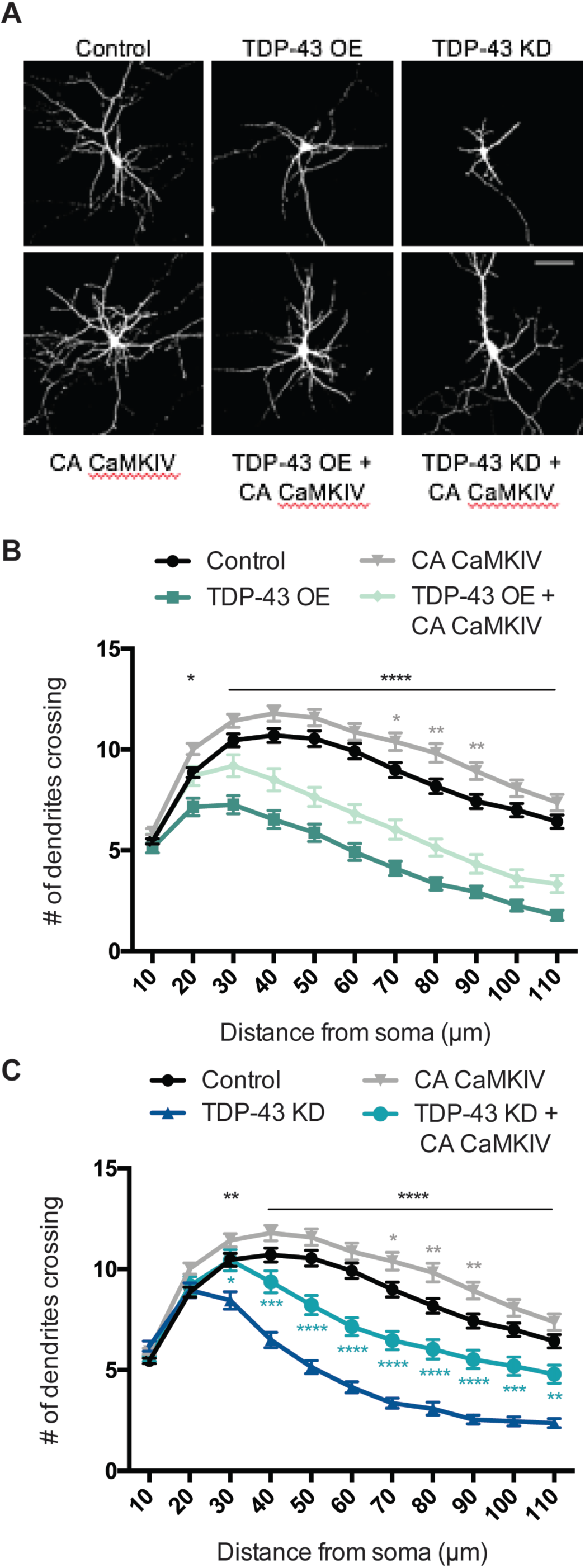
Constitutively active CaMKIV partially restores dendritic branching defects induced by TDP-43 dysfunction. **(A)** Representative images of 7 DIV cultured hippocampal neurons transfected with plasmids expressing GFP and either empty vector control, TDP-43, or an shRNA targeting TDP-43 for knockdown (upper panel). A similar set of experiments was done in neurons transfected with coexpressing CA CaMKIV (lower panel). (B) Quantification of dendritic branching via Sholl analysis for each condition (Control N = 164; TDP-43 OE = 64; CA CaMKIV = 157; TDP-43 OE + CA CAMKIV = (C) Quantification of dendritic branching via Sholl analysis for each condition. Control and CA CaMKIV traces from panel (B) re-plotted for comparison (Control, N = 164; TDP-43 KD = 82; CA CaMKIV = 157; TDP-43 KD + CA CaMKIV = 73). Two-way ANOVA with Tukey’s test. Black asterisks = control vs TDP-43 OE or TDP-43 KD; (C) turquoise asterisks = TDP-43 KD vs TDP-43 KD + CA CaMKIV; * = p < 0.05; ** = p < 0.01; *** = p < 0.001; **** = p < 0.0001. Error bars represent standard error of the mean.

As previously described our TRIBE experiment has identified potential TDP-43 targets in multiple signaling pathways culminating in CREB activation. Since our observations indicated that both loss and gain of function of TDP-43 in cultured neurons results in a decrease in CREB activation and subsequent transcriptional output (Fig.2 and 3), we sought to determine whether restoring CREB activity directly could rescue the defective dendritic morphology in both these conditions. To this end, we employed a CREB protein that was rendered constitutively active by fusing the VP16 transactivation domain from the Herpes Simplex Virus to full length CREB (CREB CA; (Tao et al. 1998). Like the experiments mentioned above with CaMKIV, hippocampal neurons were transfected at 2 DIV and fixed for morphological analysis at 5 DIV. Interestingly, the data show that expression of CREB CA in the context of both TDP-43 OE and TDP-43 KD restored dendritic complexity to near normal levels (Figure 5B-C). Expression of CREB CA alone did not affect dendritic branching, consistent with previous reports (Redmond et al. 2002). These results indicate that changes in dendritic morphology resulting from dysfunction of TDP-43 can be rescued by restoring CREB transcriptional output. Therefore, enhancing CREB signaling can be potential therapeutic avenue to address the pathological neuronal morphology that have the potential to contribute to the underlying etiology of ALS and FTD.

**Figure 5:**
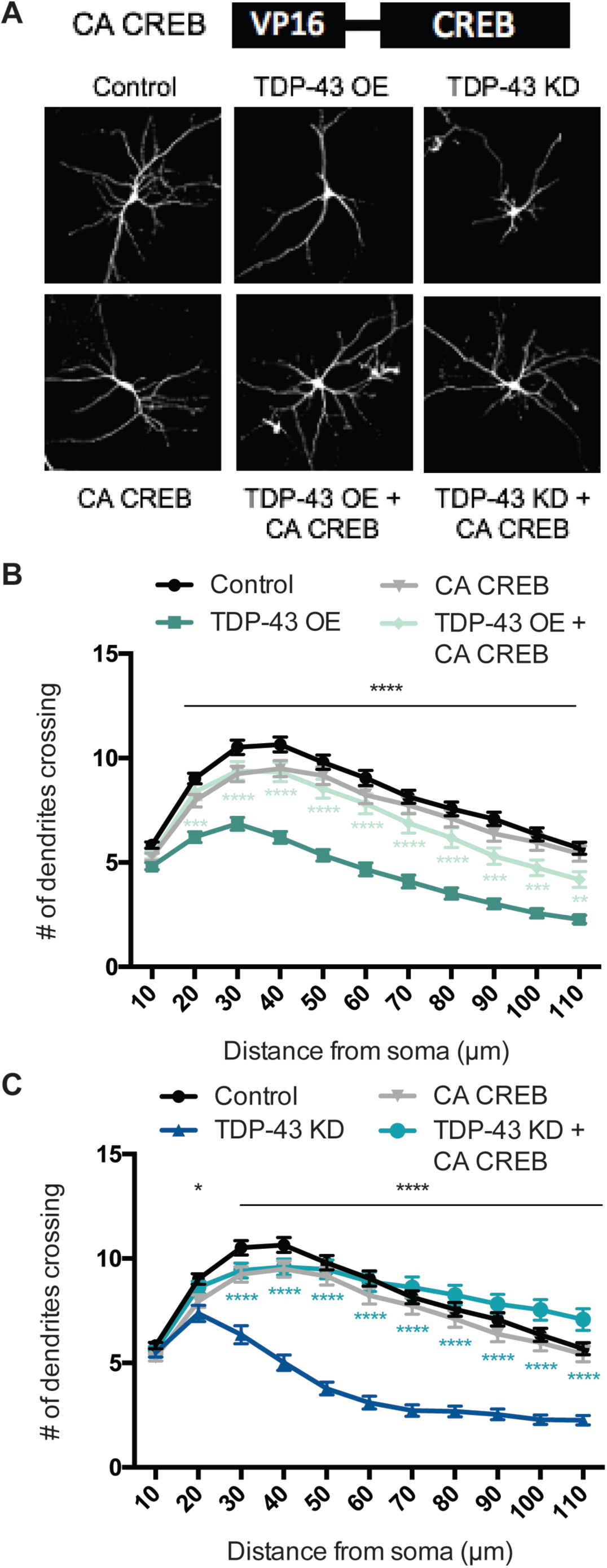
Constitutively active CREB restores dendritic branching defects induced by TDP-43 dysfunction. **(A)** Representative images of 7 DIV cultured hippocampal neurons transfected with plasmids expressing GFP and either empty vector control, TDP-43, or an shRNA targeting TDP-43 for knockdown (upper panel). A similar set of experiments was done in neurons transfected with coexpressing CA CREB (lower panel). (B) Quantification of dendritic branching via Sholl analysis for each condition (Control N = 119; TDP-43 OE = 102; CA CREB = 111; TDP-43 OE + CA CREB = 70). (C) Quantification of dendritic branching via Sholl analysis for each condition. Control and CA CREB traces from panel (B) re-plotted for comparison (Control, N = 119; TDP-43 KD = 71; CA CREB = 111; TDP-43 KD + CA CREB = 91). Two-way ANOVA with Tukey’s test. Black asterisks = control vs TDP-43 OE or TDP-43 KD; (B) light green asterisks = TDP-43 OE vs TDP-43 OE + CA CREB (C) turquoise asterisks = TDP-43 KD vs TDP-43 KD + CA CREB; * = p < 0.05; ** = p < 0.01; *** = p < 0.001; **** = p < 0.0001. Error bars represent standard error of the mean.

## Discussion

TDP-43 is a key regulator of RNA metabolism and the central component in ALS and FTD pathology. Our previous data demonstrated that TDP-43 dysfunction disrupts normal dendritic morphology, and this effect is dependent on the RNA-binding ability of TDP-43 (Herzog et al. 2017). In this paper, we applied the recently developed TRIBE method to a mammalian system and identified novel neuronal signaling pathways as targets of TDP-43. Based on the identity of the affected signaling pathways, we hypothesized and validated that CREB transcriptional activity is an important regulator of TDP-43-dependent changes in dendritic morphology. Activation of CREB alleviated the defects in dendritic morphology caused by TDP-43 dysfunction, which provides a potential druggable target for TDP-43-related neurodegenerative diseases. In addition, TRIBE was recently performed for other mammalian RBPs (Biswas et al., manuscript in preparation; H. Jin et al., manuscript in preparation), and the method has also been successfully applied to a plant system (P. Brodersen, personal communication), indicating that TRIBE will have wide applicability.

TDP-43 was reported to associate with several thousand nascent and mature RNA transcripts in neurons (Polymenidou et al. 2011; Sephton et al. 2011; Tollervey et al. 2011). One group performed TDP-43 RIP-seq with cultured cortical neurons from rats, similar to our culture system (Sephton et al. 2011), whereas the other used adult mouse brain to perform TDP-43 CLIP-seq (Polymenidou et al. 2011). The differences in source material might explain why our TDP-43 TRIBE data are more similar to the RIP-seq results (Sephton et al. 2011).

TRIBE generated a shorter list of candidate targets than both of these other studies in addition to a high level of consistency between replicates (>50% of all editing sites). The discrepancy in the number of potential targets may be due in part to the fact that the RIP and CLIP studies assayed total TDP-43 RNA including pre-mRNA. Indeed, 2/3 of the CLIP targets only have an intron signal. Although TRIBE can also identify nascent RNA targets (Menet et al. 2012; McMahon et al. 2016), we used polyA RNA in this study to specifically focus on potentially high value mRNA targets. Moreover this work does not compare directly the number of potential CLIP vs TRIBE targets, which should ideally be addressed in the same study. However, the consideration of nuclear RNA makes the number of candidate CLIP TDP-43 target mRNAs (Polymenidou et al. 2011) less than a factor of two greater than the number of candidate TRIBE TDP-43 target mRNAs identified here.

Reproducible TRIBE target genes that are also identified with other methods are less likely to be false positives and may reflect RNAs bound at high affinity by the RBP (McMahon et al. 2016; Rahman et al. 2018). In this context, TRIBE is an antibody-independent method, and RIP and CLIP targets may include false positives due to intrinsic antibody issues (Biswas et al., in preparation). The shorter TRIBE list may also reflect more false negatives, due perhaps to inefficiencies of TRIBE as previously discussed (McMahon et al. 2016). There were in addition 108 genes uniquely identified by TDP-43 TRIBE (Fig. 1D). Despite the rather small number of edited transcripts, these genes are also unlikely to be false-positives. This is because of our stringent requirement for reproducibility (the exact editing sites must be identified in both replicates), so these transcripts may therefore reflect bona fide targets that escaped identification with the other methods. Notably, some of these genes, e.g., MAPK8 and MAPK8IP1, are components of the top hit KEGG signaling pathways. In any case, the shorter candidate TRIBE list is certainly advantageous for target validation purpose.

This shorter TRIBE list also contrasts with the many thousands of target genes identified in previous TRIBE experiments with the RBP Hrp48 in *Drosophila* cultured S2 cells and in *Drosophila* neurons (Xu et al. 2018). This suggest that a general inefficiency of the TRIBE technique is unlikely to be a major explanation for the relatively small number of TDP-43 targets. Although the method could be less efficient in mammalian cells compared to *Drosophila*, Hrp48 could also be a more promiscuous RBP compared to TDP-43. Perhaps the most likely explanation is that the expression systems used in all previous *Drosophila* experiments resulted in high expression levels, likely substantial overexpression compared to endogenous protein levels and compared to the lentiviral expression of the TDP-43-hADARcd fusion protein in this study.

We and others have shown that reduced dendritic branching is a result of TDP-43 dysfunction (Majumder et al. 2012; Schwenk et al. 2016; Herzog et al. 2017). Changes in dendritic arbors have been reported in a variety of neurological disorders, and morphological changes are known to occur prior to neurodegeneration (Kulkarni and Firestein 2012; Lopez-Domenech et al. 2016; Kweon et al. 2017). Alterations in dendritic complexity may disrupt neuronal connectivity, cell-to-cell communication, and ultimately cause detrimental effects to cell health and survival. We were therefore curious whether the 532 TDP-43 targets identified by TRIBE functioned in specific signaling pathways that might impact dendritic morphology. CREB is an activity-dependent transcription factor with a well-established role in regulating dendritic complexity (Tan et al. 1996; Tao et al. 1998; Watson et al. 2001; Redmond et al. 2002; Alboni et al. 2011) and DAVID GO term and KEGG pathway analysis revealed that the TDP-43 TRIBE target gene list was significantly enriched for signaling pathways that converge on CREB activation. Importantly, CREB signaling and gene expression have previously been shown to be downregulated in several neurodegenerative diseases (Mantamadiotis et al. 2002; Saura and Valero 2011). Taken together with the 3’UTR preference of the TDP-43 TRIBE signals, our data suggest that TDP-43 post-transcriptional regulation of discrete targets is upstream of CREB function, which contributes to the dendritic branching defects of the pathologic state.

Cytoplasmic TDP-43 aggregation may deplete functional cytoplasmic as well as nuclear TDP-43, thereby decreasing the function of this protein in processes like translation as well as nuclear events like RNA splicing. It would also not be surprising if ALS-like phenotypes in TDP-43 aggregating neurons were due to synergy from the decrease in TDP-43 activity in both compartments. Moreover, proper cytoplasmic function may require that TDP-43 loads onto pre-mRNA in the nucleus and is then exported to the cytoplasm associated with its mRNA targets. From this perspective, TDP-43 cytoplasmic function(s) may not be completely distinct from its nuclear function(s).

Signal transduction pathways controlled by stimuli such as neurotrophic growth factors and Ca^2+^ have been previously implicated in a subset of ALS patient etiology (Henriques et al. 2010; Leal and Gomes 2015). Interestingly, the gene encoding the pore-forming subunit of the N-type voltage-gated Ca^2+^ channel, *Cacna1B*, was commonly identified as a TDP-43 RNA target by all 3 RNA target identification methods. This example suggests that Ca^2+^ signaling events both upstream and downstream of Ca^2+^ entry may be disrupted upon TDP-43 dysfunction. Importantly, Ca^2+^ signaling also converges on CREB activation.

Our data demonstrate that co-expression of CA CREB with TDP-43 OE or TDP-43 KD fully rescued dendritic branching. In a complementary pathway, loss of TDP-43 in primary hippocampal neurons reduces surface expression of the growth factor receptor ErbB4 and impairs downstream signaling, also resulting in decreased dendritic complexity (Schwenk et al. 2016). In addition, dendritic branching can be restored in TDP-43 depleted neurons by overexpressing ErbB4 (Schwenk et al. 2016). These data and ours taken together suggest that multiple signaling pathways that influence dendritic branching are affected by TDP-43 dysfunction.

Interestingly, our GO term analyses also identified enriched processes that also establish neuronal communication at chemical synapses: synapse assembly and dendritic spine morphogenesis (Fig. 1). It is therefore possible that TDP-43 dysfunction disrupts dendritic morphogenesis processes directly, as well as indirectly through signaling pathways that are engaged by synaptic transmission and in turn impact CREB activation. This may explain why CaMKIV CA expression can only partially rescue the dendritic complexity phenotype.

In conclusion, our findings reveal a role for TDP-43 in the regulation of dendritic morphology via RNA targets that converge on dendritic branching. Given that reduced dendritic complexity may precede neurodegeneration (Lopez-Domenech et al. 2016), a delay between the change in morphology and neuronal function serves as a potential window for therapeutic intervention. Although many early ALS studies focused on the morphological changes that happen at the distal axon in the peripheral nervous system (Chou and Norris 1993), our data support a model that alterations of the dendritic arbor in the central nervous system may also contribute to disease pathogenesis. Indeed, ALS patients display reduced dendrite morphologies in both upper and lower motor neurons which may then encourage neurodegeneration (Takeda et al. 2014; Genc et al. 2017). Our work therefore encourages more effort to reestablish nervous system connectivity as a therapeutic target in ALS/FTD.

## Methods

The datasets generated during and/or analysed during the current study are available from the corresponding author on reasonable request. All animal procedures were approved by the Brandeis University Institutional Animal Care and Usage Committee and all experiments were performed in accordance with relevant guidelines and regulations.

### Plasmids cloning

pCMV vector was used to create pCMV-TRIBE constructs for HEK293T cell expression. hTDP-43-hADARcd-E488Q, hTDP-43-5FL-hADARcd-E488Q or hADARcd-E488Q was inserted into the vector by Gibson Assembly. pCMV-TRIBE plasmids were individually transfected into HEK293T cells by standard calcium phosphate transfection. pCMV-EGFP was co-transfected to monitor transfection efficiency.

pLV-hSyn-RFP was a gift from Edward Callaway (Salk Institute for Biological Studies, La Jolla, CA) (Addgene plasmid # 22909). pLV-hSyn-RFP plasmid was linearized by cutting it with BamHI. A myc-hTDP-43-P2A cassette with complementary overhangs was synthesized using BioXP Custom cloning (Synthetic Genomics Inc.) and cloned in frame by Gibson Assembly. For TDP-43 TRIBE plasmids cloning, pLV-hSyn-RFP were cut with PmeI and AgeI. hTDP-43-hADARcd-E488Q or hADARcd-E488Q fragment with complementary overhang was inserted into the digested backbone by Gibson Assembly. RFP remains in frame and serves as a marker for transfection efficiency.

### Neuronal Culture and Transfection

12 mm glass coverslips were coated with poly-D-lysine (20 μg/ml) and laminin (3.4 μg/ml) in 24 well plates for 1-24 hours. Then the coverslips were washed 3x with deionized H_2_O and 2x with Neurobasal Medium. Dissociated hippocampal neurons from E18 rat embryos were cultured on an astrocyte feeder layer, as described previously (Ghiretti and Paradis 2011), at a density of 80,000 neurons/well (for Sholl Analysis). Luciferase assays were performed with dissociated cortical neurons from E18 rat embryos without an astrocyte feeder layer. Cultured cortical neurons were plated at a density of 500,000 neurons/well. Both hippocampal and cortical cultures were grown in Neurobasal Medium supplemented with B27 (Thermo Fisher) at 37°C.

For Sholl Analysis, neurons were transfected on DIV 2 by the calcium phosphate method (Xia et al. 1996), which transfects a low percentage of cells allowing the genetically manipulated neuron to grow in an environment surrounded by unaffected or wild type cells. This method is also ideal to visualize the dendritic morphology of individual cells in a dense population. For all experimental conditions, unless noted otherwise, neurons were co-transfected with pCMV-GFP plasmid at 500 ng/well to visualize neuronal morphology. Along with pCMV-GFP, neurons were either transfected with pCMV-TDP-43 (TDP-43 OE) at 500 ng/well or shRNAs expressed from pSuper targeting endogenous TDP-43 at 33 ng/well (TDP-43 KD). For coexpression experiments with constitutive active forms of CaMKIV and CREB, plasmids expressing these genes were transfected at 100 ng/well wither alone or with TDP-43 OE or TDP-43 KD.

For luciferase assays, neurons were transfected on DIV 2 by the calcium phosphate method with either an empty vector control, TDP-43 OE, or TDP-43 KD at the same concentrations mentioned above. In addition, each condition was cotransfected with reporter constructs containing the firefly luciferase gene downstream of concatemerized CREB binding sites at 500 ng/well. A separate reporter construct expressing *Renilla* luciferase being driven by the thymidine kinase promoter or the EF-1α promoter was transfected at 100 ng/well to serve as an internal control for normalizing transfection efficiency. On DIV 6, neurons were treated with 1 μM TTX to block sodium channels. On DIV 7, 55 mM KCl was added to a subset of neurons from each condition. After 6 hours of KCl treatment, neurons were washed with 1X PBS and lysed with passive lysis buffer (Promega) and luciferase levels were measured using a dual luciferase assay. Firefly luciferase was measured by adding 100 μl of 75 mM HEPES, pH 8.0, 5 mM MgSO_4_, 20 mM DTT, 100 mM EDTA and 530 μM ATP, 0.5 mM d-luciferin and 0.5 mM coenzyme A. *Renilla* luciferase was measured using 100 μl of 25 mM Na_4_PPi, 10 mM NaOAc, 15 mM EDTA, 0.5 M Na_2_SO_4_, 1 M NaCl, and 0.1 mM Coelenterazine, pH 5.0. All conditions were repeated at least three times in triplicate and normalized to *Renilla* luciferase and then to the untreated control condition.

For TDP-43 TRIBE experiments, dissociated cortical neurons from E18 rat embryos were cultured in a 6 well plate at 4 Million neurons/well and grown in glia conditioned Neurobasal Medium supplemented with B27 at 37°C. Neurons were transduced at DIV 5 with TDP-43 TRIBE virus (packaged by Vigene Biosciences, Inc.) and RNA was harvested 3 days later on DIV 7. (Rodriguez et al. 2012; McMahon et al. 2016; Rahman et al. 2018).

### RNA-seq library preparation and sequencing

Transfected HEK293T cells or neurons were harvested 3 days after transfection in TRIzolTM reagent (Thermo Fisher Scientific, 15596026). RNA was extracted as recommended by TRIzol product manual, dissolved in Milli-Q H2O and stored in −80°C freezer. 1-2µg of total RNA was poly(A) beads selected and used to prepare for mRNA-seq library with NEXTFLEX^®^ Rapid Directional qRNA-Seq Kit. The library product was quantified with Agilent 4200 TapeStation System and normalized to 2nM for sequencing purpose. Libraries were pooled and then pair-end sequenced for 75bp by NextSeq 500 System by Illumina^®^ (NextSeq^®^ High Output Kit v2, 75 cycles). Each libraries had ∼50 million pair-end reads.

### RNA-seq data analysis

RNA sequencing data were analyzed as described in Rahman et al. 2018 and scripts used are available on GitHub (https://github.com/rosbashlab/HyperTRIBE). Hg38 genome was used for HEK293T cell data and Rn6 genome was used for rat neuron data. PCR duplication was removed using UMIs included in the NEXTFLEX kit. Sequencing data of RNA from wild type HEK293T cells or untransfected rat neurons was used as the reference for calling RNA editing events, respectively. The criteria for RNA editing events were: 1) The nucleotide is covered by a minimum of 11 reads in each replicate; 2) More than 80% of wild type RNA reads at this nucleotide is A with zero G (use the reverse complement if annotated gene is in the reverse strand); 3) A minimum of 5% G is observed at this site in mRNA (or C for the reverse strand); 4) Editing sites are present in both replicates. Distribution of editing sites in each mRNA region was determined using bedtools, intersecting editing sites with RefSeq annotated CDS, 5’UTR and 3’UTR. Distribution of the transcriptome in these regions was calculated using RSeQC (http://rseqc.sourceforge.net/).

Gene expression profiles were generated by feeding bam files into cufflinks, and ranked by FPKM (Fragments Per Kilobase of transcript per Million mapped reads) value. Top 3000 genes were used as the background list for DAVID analysis.

### Immunostaining

Following primary antibodies were used for immunostaining, Anti-Phospho-CREB-Ser133 (87G3) (Rabbit mAb # 9198) from Cell Signaling Technologies (used at 1:500), Anti-CREB (Cat. No. 06-863) from Millipore Sigma (used at 1:100), Anti-CREST (12439-1-AP) from Proteintech (used at 1:100), Anti-TFIIS, (611204) from BD Transduction Laboratories (used at 1:275). Secondary antibodies were conjugated to Cy-3 or Cy-5 (1:500, Jackson ImmunoResearch Laboratories).

Coverslips containing transfected neurons were fixed as described above followed by 3 washes with 1X PBS. The coverslips were then incubated with primary antibodies) diluted in gelatin blocking buffer at 4°C overnight in a humidified chamber. Coverslips were then washed 3x with 1X PBS and incubated with secondary antibodies at room temperature for 2 hours. Coverslips were then washed 3x with 1X PBS, submerged in MilliQ H_2_O and then mounted on glass microscope slides with Aquamount (Lerner Laboratories). Neurons were imaged using a spinning-disk confocal system consisting of a Nikon Ni-E upright microscope, a Yokogawa CSU-W1 spinning-disk head, and an Andor iXon 897U electron-multiplying charge-coupled device camera using 40× (numerical aperture [NA] 1.3) or 60× (NA 1.4) oil immersion objectives. Identical acquisition settings were used for laser power, exposure time and detector gain across conditions, and data were acquired and analyzed in a blinded manner.

### Image Analysis

To quantify pCREB levels, sum projections were generated from individual Z stacks using ImageJ. In ImageJ, using GFP as a marker for transfected cells, a region of interest was drawn around the soma and fluorescence intensity was measured in that area in the pCREB channel. Then, TFIIS fluorescence intensity was measured using the same region of interest. pCREB intensity was normalized to TFIIS intensity for every cell. The same procedure was used for the CREB antibody.

### Analysis of Neuronal Morphology

Transfected neurons were fixed at DIV 7 with 4% paraformaldehyde + 4% sucrose solution in PBS for 8 minutes at room temperature and washed 3x with 1X PBS. An Olympus Fluoview 300 confocal microscope using a 20X oil objective (numerical aperture [NA] 0.85) was used to image GFP-transfected cells (5-10 steps at 1 μm optical sectioning). All images were analyzed in ImageJ using Sholl Analysis where Maximum Intensity Projections (MIP) were generated from individual Z stacks. To quantify dendritic complexity, 11 concentric circles (10 μm intervals) centered at the soma were overlaid on the image and the number of dendritic crossings was counted for each 10 µm interval. For each experimental condition, a total of 10-30 neurons were analyzed from 2-3 coverslips and each experiment was independently repeated at least twice as a biological replicate. Imaging and analysis were performed in a blinded manner, and unblinded only after analysis.

## Acknowledgements

This work was supported by a Blazeman Foundation for ALS Research Postdoctoral Fellowship (M.D.) and by NIH grants R01DA037721 (M.R.), R01AG052465 (M.R.), and R01NS065856 (S.P.).

---

**Table S1:**
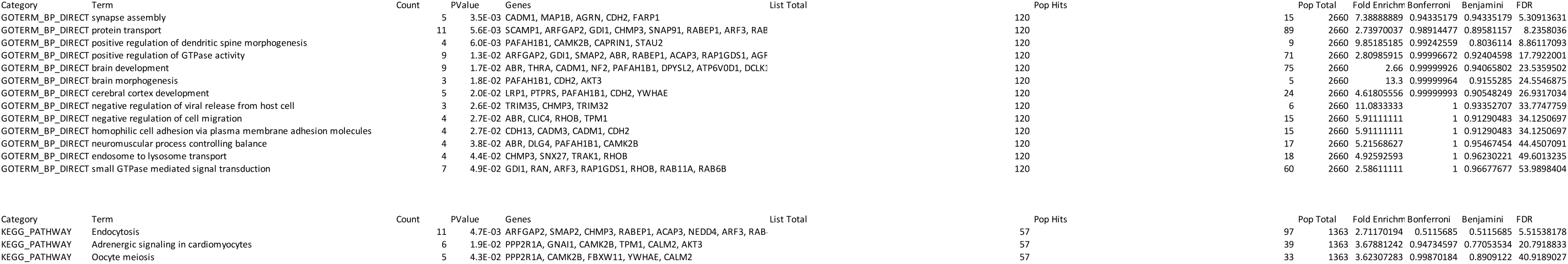
Overlapping Genes from all 3 methods.

**Table S1:** Genes from TDP-43 RIP.

| Category | Term | Count | PValue | Genes | List Total | Pop Hits | Pop Total | Fold Enrichm | Bonferroni | Benjamini | FDR |
| --- | --- | --- | --- | --- | --- | --- | --- | --- | --- | --- | --- |
| GOTERM_BF | in utero embryonic development | 46 | 3.6E-03 | NM_001135923, NM_001034068, NM_001034069, NM_024: | 1444 | 63 | 2660 | 1.34502924 | 0.99999985 | 0.99999985 | 6.57224945 |
| GOTERM_BF | cell adhesion | 30 | 4.0E-03 | NM_012916, NM_001107740, NM_001033970, NM_031333, | 1444 | 38 | 2660 | 1.45429363 | 0.99999998 | 0.99984194 | 7.29708764 |
| GOTERM_BF | negative regulation of gene expression | 36 | 5.9E-03 | NM_001013059, NM_017041, NM_001077585, NM_022681, | 1444 | 48 | 2660 | 1.38157895 | 1 | 0.99980248 | 10.4847878 |
| GOTERM_BF | negative regulation of cell growth | 23 | 1.4E-02 | NM_021660, NM_001109541, NM_019197, NM_134368, Nv | 1444 | 29 | 2660 | 1.46098004 | 1 | 0.99999966 | 22.719452 |
| GOTERM_BF | RNA secondary structure unwinding | 15 | 1.6E-02 | NM_001015018, NM_001015005, NM_001106793, NM_0010 | 1444 | 17 | 2660 | 1.625387 | 1 | 0.99999879 | 25.5354089 |
| GOTERM_BF | protein complex assembly | 12 | 2.4E-02 | NM_057140, NM_022210, NM_001109880, NM_019621, Nv | 1444 | 13 | 2660 | 1.70040486 | 1 | 0.99999996 | 35.9697558 |
| GOTERM_BF | synapse assembly | 13 | 3.7E-02 | NM_031606, NM_134336, NM_001107287, NM_053992, Nv | 1444 | 15 | 2660 | 1.59649123 | 1 | 1 | 50.7306656 |
| GOTERM_BF | cell cycle | 25 | 3.8E-02 | NM_133589, NM_013145, NM_052981, NM_001005543, Nv | 1444 | 34 | 2660 | 1.35448916 | 1 | 1 | 51.8767129 |
| GOTERM_BF | cell-cell adhesion | 56 | 4.1E-02 | NM_024397, NM_031986, NM_001009692, NM_001025410, | 1444 | 86 | 2660 | 1.1995104 | 1 | 1 | 54.6509767 |
| GOTERM_BF | neuron differentiation | 29 | 4.7E-02 | NM_001009674, NM_153297, NM_001108596, NM_024374, | 1444 | 41 | 2660 | 1.3029525 | 1 | 1 | 59.3970683 |
| GOTERM_BF | exocytosis | 16 | 4.8E-02 | NM_130412, NM_001025686, NM_031693, NM_057097, Nv | 1444 | 20 | 2660 | 1.47368421 | 1 | 1 | 59.9349669 |
| KEGG_PATH | Synaptic vesicle cycle | 29 | 1.1E-04 | NM_001011927, NM_212490, NM_001107511, NM_053788, | 725 | 33 | 1363 | 1.65212121 | 0.02929142 | 0.02929142 | 0.14886351 |
| KEGG_PATH | Phosphatidylinositol signaling system | 18 | 8.5E-04 | NM_031606, NM_021660, NM_053316, NM_022301, NM_00 | 725 | 19 | 1363 | 1.78105263 | 0.19893737 | 0.10497898 | 1.1053846 |
| KEGG_PATH | Endocrine and other factor-regulated calcium reabsorption | 16 | 7.6E-03 | NM_013113, NM_012913, NM_022952, NM_001100922, Nv | 725 | 18 | 1363 | 1.67111111 | 0.86260106 | 0.48398643 | 9.46768617 |
| KEGG_PATH | Cell adhesion molecules (CAMs) | 13 | 1.2E-02 | NM_017022, NM_001007728, NM_134336, NM_017345, Nv | 725 | 14 | 1363 | 1.74571429 | 0.95307161 | 0.53456513 | 14.2123995 |
| KEGG_PATH | mRNA surveillance pathway | 29 | 1.3E-02 | NM_001008295, NM_001011890, NM_001106785, NM_147: | 725 | 39 | 1363 | 1.39794872 | 0.96287267 | 0.48246618 | 15.2136101 |
| KEGG_PATH | Prostate cancer | 19 | 2.3E-02 | NM_031606, NM_031515, NM_171992, NM_080766, NM_1: | 725 | 24 | 1363 | 1.48833333 | 0.9977502 | 0.63801486 | 26.3260885 |
| KEGG_PATH | Rheumatoid arthritis | 15 | 2.8E-02 | NM_001011927, NM_031604, NM_212490, NM_199386, Nv | 725 | 18 | 1363 | 1.56666667 | 0.99941831 | 0.65500505 | 31.1544539 |
| KEGG_PATH | Adrenergic signaling in cardiomyocytes | 28 | 2.9E-02 | NM_012913, NM_013145, NM_001034068, NM_001034069, | 725 | 39 | 1363 | 1.34974359 | 0.99948697 | 0.61205691 | 31.5863828 |
| KEGG_PATH | Inflammatory mediator regulation of TRP channels | 13 | 3.0E-02 | NM_031977, NM_001100922, NM_022498, NM_013065, Nv | 725 | 15 | 1363 | 1.62933333 | 0.99966758 | 0.58930392 | 33.0579968 |

**Table S1:**
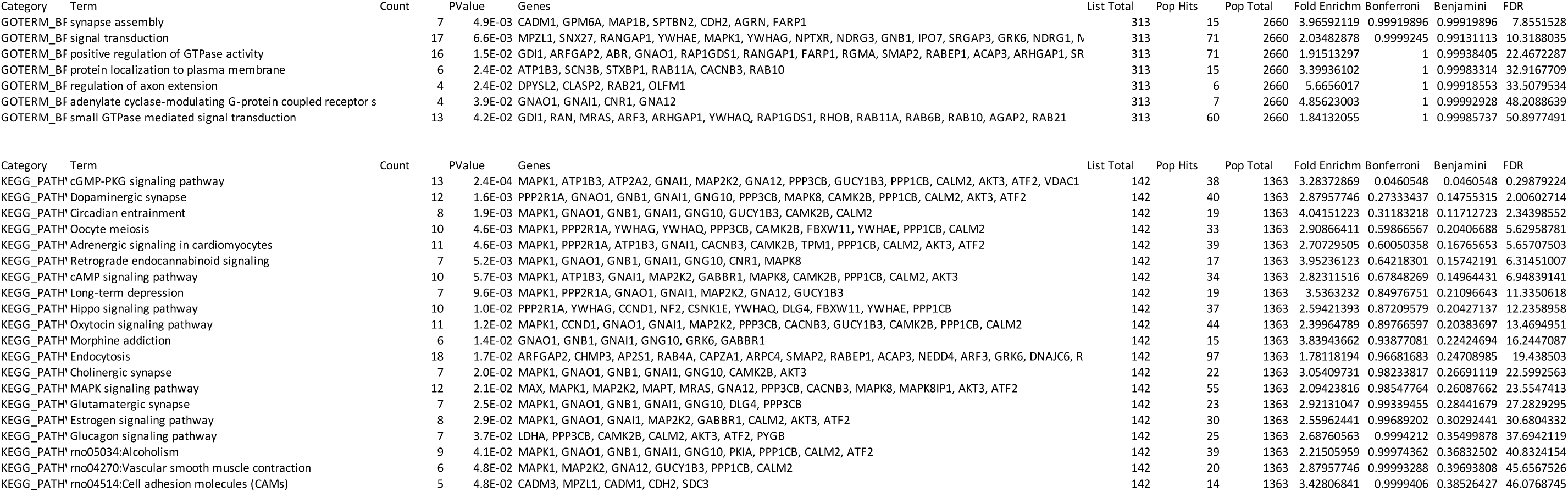
Genes from TDP-43 TRIBE.

